# The capsid precursor protein of astrovirus VA1 is proteolytically processed intracellularly

**DOI:** 10.1101/2022.04.28.489973

**Authors:** Catalina Aguilera-Flores, Tomás López, Fernando Zamudio, Carlos Sandoval-Jaime, Edmundo I. Pérezc, Susana López, Rebecca DuBois, Carlos F. Arias

**Affiliations:** Departamento de Genética del Desarrollo y Fisiología Molecular, Universidad Nacional Autónoma de México, Cuernavaca, Morelos, Mexico; Departamento de Medicina Molecular y Bioprocesos, Instituto de Biotecnología, Universidad Nacional Autónoma de México, Cuernavaca, Morelos, Mexico; Department of Biomolecular Engineering, University of California Santa Cruz, Santa Cruz, California, USA

**Author notes:** Corresponding author: Carlos F. Arias.

**Keywords:** VA1 astrovirus, proteolytical cleavage, RNA virus, capsid processing

## Abstract

Human astrovirus VA1 has been associated with neurological disease in immunocompromised patients, and its recent propagation in cell culture has opened the possibility to study its biology. Unlike classical human astroviruses, VA1 growth was found to be independent of trypsin during virus replication *in vitro*. In this work, we show that despite its independence on trypsin activation for cell infection, the VA1 capsid precursor protein, of 86 kDa (VP86), is processed intracellularly, and this proteolytic processing is important for astrovirus VA1 infectivity. Antibodies raised against different regions of the capsid precursor showed that the polyprotein can be processed starting at either its amino-or carboxy-terminal end, and they allowed us to identify that proteins of about 33 (VP33) and 38 (VP38) kDa constitute the core and the spike proteins of the mature infectious virus particles, respectively. The amino-terminal end of the spike protein was found to be Thr-348. Whether the protease involved in intracellular cleavage of the capsid precursor is of viral or cellular origin remains to be determined, but the cleavage is independent of caspases. Also, trypsin is able to degrade the capsid precursor but has no effect on VP34 and VP38 proteins when assembled into virus particles. These studies provide the basis for advancement of the knowledge of astrovirus VA1 cell entry and replication.

**IMPORTANCE:** Human astrovirus VA1 has been associated with neurological disease in immunocompromised patients. Its recent propagation in cell culture has facilitated the study of its biology. In this work, we show that despite the ability of this virus to grow in the absence of trypsin, a marked feature of human classical astroviruses, the capsid precursor protein of astrovirus VA1 is cleaved intracellularly to yield the mature infectious particles, formed by two polypeptides, VP33 that constitutes the core domain of the virus particle, and V38 that forms the spike of the virus. These studies provide a platform to advance our knowledge on astrovirus VA1 cell entry and replication.

## INTRODUCTION

Human astrovirus (HAstV) is an important etiological agent of gastroenteritis, affecting mainly children, the immunocompromised, and the elderly (1, 2). Most of these infections are associated with the so-called canonical or classic HAstVs, which comprise eight different serotypes (HAstV-1 to -8) (1). In recent years, highly divergent MLB and VA genotypes of human astroviruses have been detected in stool samples, although their association with gastroenteritis has not been clearly demonstrated yet (3, 4). They have also been reported as a cause of meningitis and encephalitis in immunosuppressed patients, increasing the interest in the study of these viruses (3). The most commonly associated astrovirus genotype in human neurological disease is VA1. A recent report on the seroprevalence of neutralizing antibodies from two cohorts of adult and pediatric serum samples found the seropositivity rate in adults to be approximately 77%, and it was concluded that a majority of humans are exposed to VA1 between 2 and 9 years of age (5).

HAstVs are small, nonenveloped viruses with a single-stranded, positive-sense RNA genome of about 6.8 kb that comprises three open reading frames (ORFs). ORFs 1a and 1b encode the virus non-structural proteins, including a VPg, a viral protease, and the RNA-dependent RNA polymerase, while ORF2 codes for the virus capsid precursor protein (1, 6). Viral and cellular proteases are involved in processing polyproteins nsp1a and nsp1ab to produce the functional nonstructural proteins of the virus (6, 7). On the other hand, the capsid precursor structural polyprotein (VP90) of HAstV-8 self-assembles intracellularly and is processed by caspases to produce a virus particle formed by a 70-kDa protein (VP70) (7). This immature particle egresses from the cell and is further processed by extracellular proteases; *in vitro*, the treatment of this particle with trypsin yields a mature infectious virus formed by two protein species, VP34 and VP27 (7, 8). VP34 constitutes the shell of the virus particle (core domain) and VP27 constitutes the 30 dimeric globular spikes that protrude from the virion (spike domain) (7-9). Trypsin treatment has been shown to increase the infectivity of HAstV 10^3^ to 10^5^-fold (10-12).

Human astrovirus VA1 has been adapted to grow in cell culture, and it has been shown that its infectivity does not depend on trypsin treatment (13). However, it has not been determined if the ORF2 precursor protein is proteolytically cleaved during virus replication and its possible role on infection. In this work, we show that the VA1 capsid precursor protein, of approximately 86 kDa (VP86), is processed intracellularly through a collection of intermediate cleavage products to generate final products of about 33 (VP33) and 38 (VP38) kDa. These proteins form, respectively, the core and spike domains, of mature infectious virus particles. We also show that caspases and trypsin are not involved in the proteolytically processing pathway of VP86 and that the cleavage of this precursor is important for cell infection.

## RESULTS

### The VA1 capsid precursor protein is proteolytically processed

It has been shown that VA1 does not require trypsin treatment to activate its infectivity, and in concordance with this observation, the virus was reported to grow in the presence of 10% FCS (13). To evaluate the effect of FCS on the production of the VA1 capsid protein, Caco-2 cells were infected with trypsin-untreated virus at a multiplicity of infection (MOI) of 3, and were incubated for various times, and up to 120 hours post-infection (hpi). At the indicated times the cell media and the cells were collected independently, and the synthesis of viral proteins was assessed by Western blot analysis. For this, a rabbit hyperimmune serum (antibody DB) raised against a recombinant protein encoding the predicted core domain and subsequent linker region of the VA1 capsid protein was used (see Material and Methods). We found no major differences in the kinetics of intracellular viral protein synthesis in the presence or absence of 10% FCS, but most surprisingly was the observation that the capsid precursor protein of approximately 86 kDa was processed into smaller polypeptides between 30 and 40 kDa under both conditions (Figs. 1A and 1B). The fact that processing occurs in the absence of trypsin in the cell medium, and also in the presence of FCS in the extracellular medium, suggests that the cleavage occurs intracellularly. We started to detect viral proteins at 24 hpi, and their synthesis seemed to reach a plateau at 48 hpi (Fig. 1). On the other hand, extracellular viral proteins, present in the cell medium, were initially detected at 72 hpi, although the time of detection varied from experiment to experiment. The capsid protein in the media was also found to be also processed, showing a cleavage pattern similar to that observed for cell-associated viral proteins (Fig. 1C). Of relevance, at least 6 protein bands were recognized, which are presumably proteolytic products of the capsid precursor protein, with apparent molecular weights ranging from 33 to 86 kDa. The number of these bands varied in different experiments, suggesting that some of them are intermediate cleavage products that most likely render the proteins that constitute the mature virus.

**Fig. 1.**
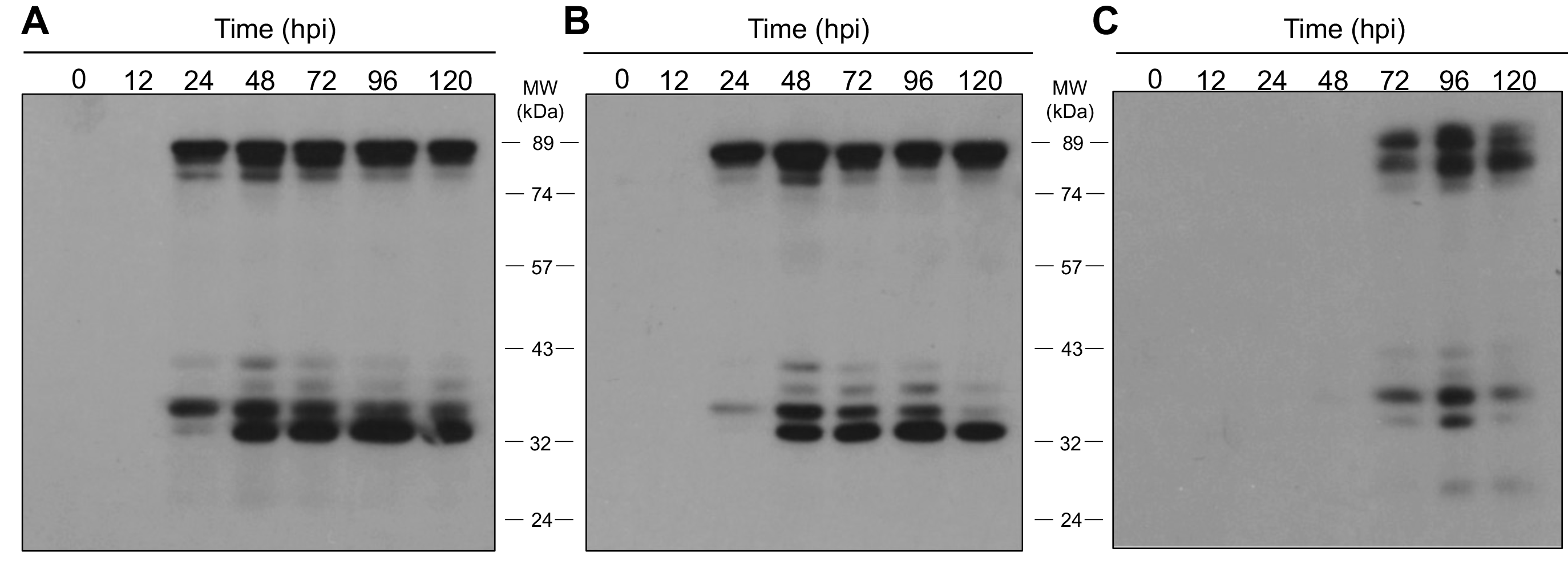
The capsid precursor protein of astrovirus VA1 is proteolytically processed. Monolayers of Caco-2 cells were infected with VA1 at an MOI of 3 in the presence (A) or absence (B) of 10% FCS in DMEM-HG, and the infection was left to proceed for the indicated times post infection (hpi) at 37°C. At the end of the incubation periods the cell media and the cell monolayers were collected independently; to this end, the medium was collected and the cells were washed twice with DMEM-HG and lysed by three cycles of freeze-thawing. Laemmli sample buffer was added to samples and they were analyzed by Western blot using a rabbit hyperimmune serum (antibody BD). (C) Western blot analysis with antibody DB of the viral proteins present in the culture media of the Caco-2 cells infected with VA1 in the absence of FCS, at the indicated hpi. The MW markers are indicated.

### Single-cycle replication cycle of VA1

A single-cycle growth curve was performed in the absence of serum to characterize the replication kinetics of VA1 in Caco-2 cells. For this, cells were infected with a MOI of 5 and, at 0 to 72 hpi the extracellular medium and the cells were independently collected and the amount of infectious virus was determined in each fraction (Fig. 2). We observed an eclipse period of approximately 12 h, followed by a burst phase from 12 to 48 hpi, in which infectious viral particles are produced and accumulate inside the cells. The stationary phase started at 48 hpi, a stage in which virus production markedly slows. On the other hand, the virus was detected in the extracellular medium starting at 18 hpi, and increased gradually until the released particles reached about 10% of the total infectious virus produced at 72 hpi.

**Fig. 2.**
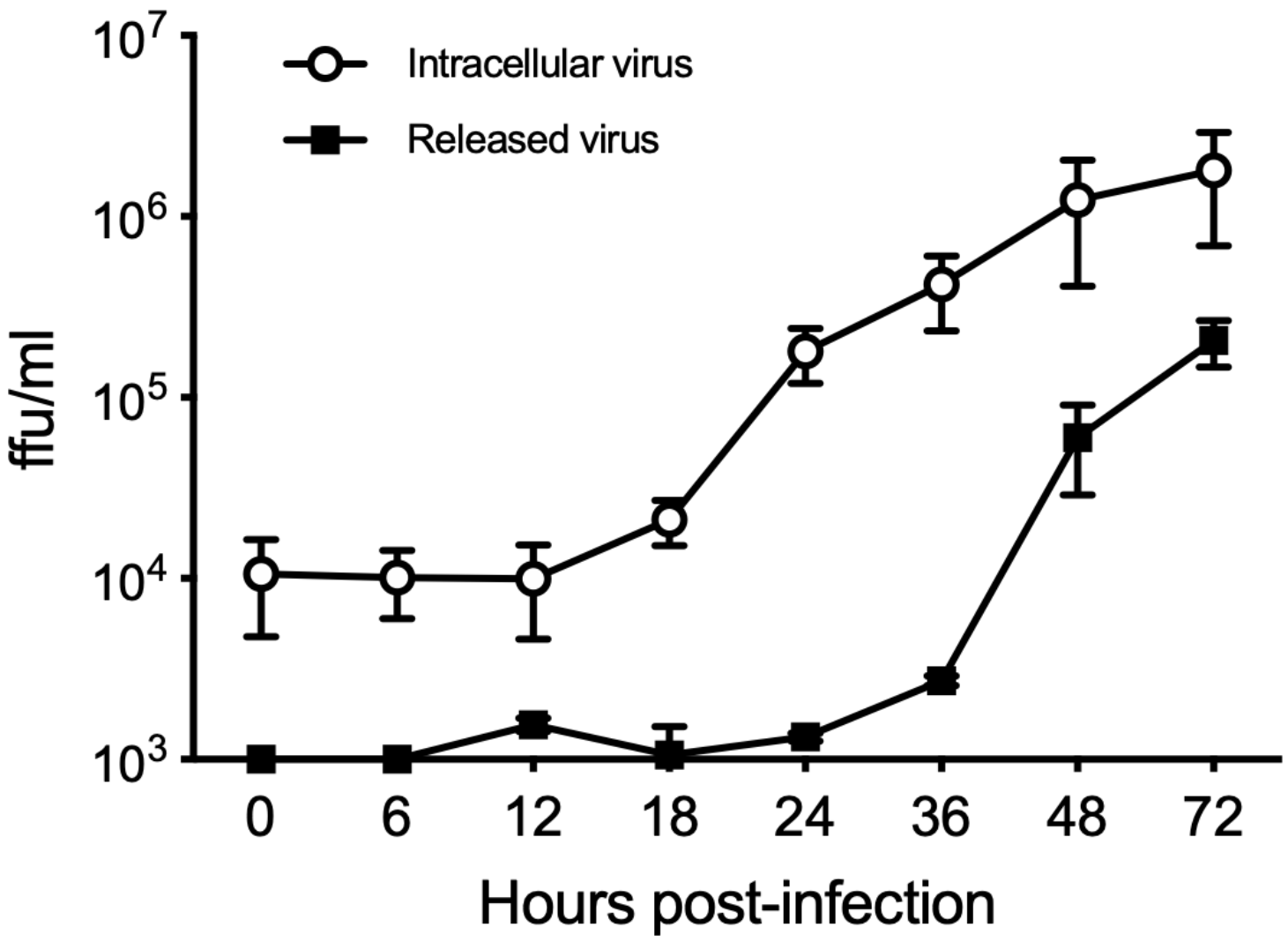
Single-cycle replication curve of astrovirus VA1. Monolayers of Caco-2 cells were infected with VA1 at an MOI of 5 in DMEM-HG, and the infection was left to proceed for the indicated times post infection (hpi) at 37°C. At the end of the incubation periods the cell media and the cell monolayers were collected independently; to this end, the medium was removed and the cells were washed twice with DMEM-HG, additional medium was added, and the cells were lysed by three cycles of freeze-thawing. The virus titer in both fractions was determined by an immunoperoxidase focus-forming assay as described in Materials and Methods. Data represent the mean±SD of three independent experiments.

### Identification of the proteins that compose the mature infectious virus

To characterize the capsid precursor and proteolytic products encoded by VA1 ORF2, rabbit polyclonal antibodies were prepared against two recombinant proteins produced in *E. coli* and against two synthetic peptides. These immunogens were chosen to generate antibodies that specifically recognize the core domain and subsequent linker region (antibody DB), a smaller fragment of the core (α-core), spike (α-spike), and carboxyl-terminal (α-COOH) regions of the precursor protein, based on what is known of these regions in the capsid protein precursor of classical astroviruses (7) (See Material and Methods, and Fig. 3A). These antibodies were used to analyze by Western blot a preparation of purified VA1 virus and a lysate of VA1-infected Caco-2 cells harvested at 48 hpi. In the infected cell lysate, the DB antibody recognized a major precursor protein of 86 kDa and a minor second precursor of about 80 kDa. Four additional proteins, of 33, 36, 38, and 42 kDa, were also recognized (Fig. 3B). On the other hand, when the purified virus was characterized with this hyperimmune serum, two major proteins were detected, of 33, and 38 kDa, suggesting that the DB antibody probably recognized regions in both the core protein and in the N-terminus of the spike protein (see below for 38 kD being the spike protein). A faint third protein of 35 kDa was also observed (Fig. 3B). When the polyclonal antibody against the core (α-core) was used, we found that not only it recognized the virus core domain, but also the 86-kDa precursor protein and three proteins of 33, 35 and 42 kDa in the infected cell lysate, while in the purified virus preparation a single protein of 33 kDa was observed (Fig. 3C). These results indicate that the 33 kDa band corresponds to the mature core protein of the virus, while the proteins of 35 and 42 kDa appear to be processing intermediates of the final core protein assembled in the virus particle. In different experiments the precursor protein of 35 kDa showed an apparent molecular weight that ranged between 34.5 to 36.2 kDa, while the 42-kDa ranged between 40.5 and 42.6 kDa, suggesting that alternative cleavage sites can lead to the mature proteins assembled in the virus particles.

**Fig. 3.**
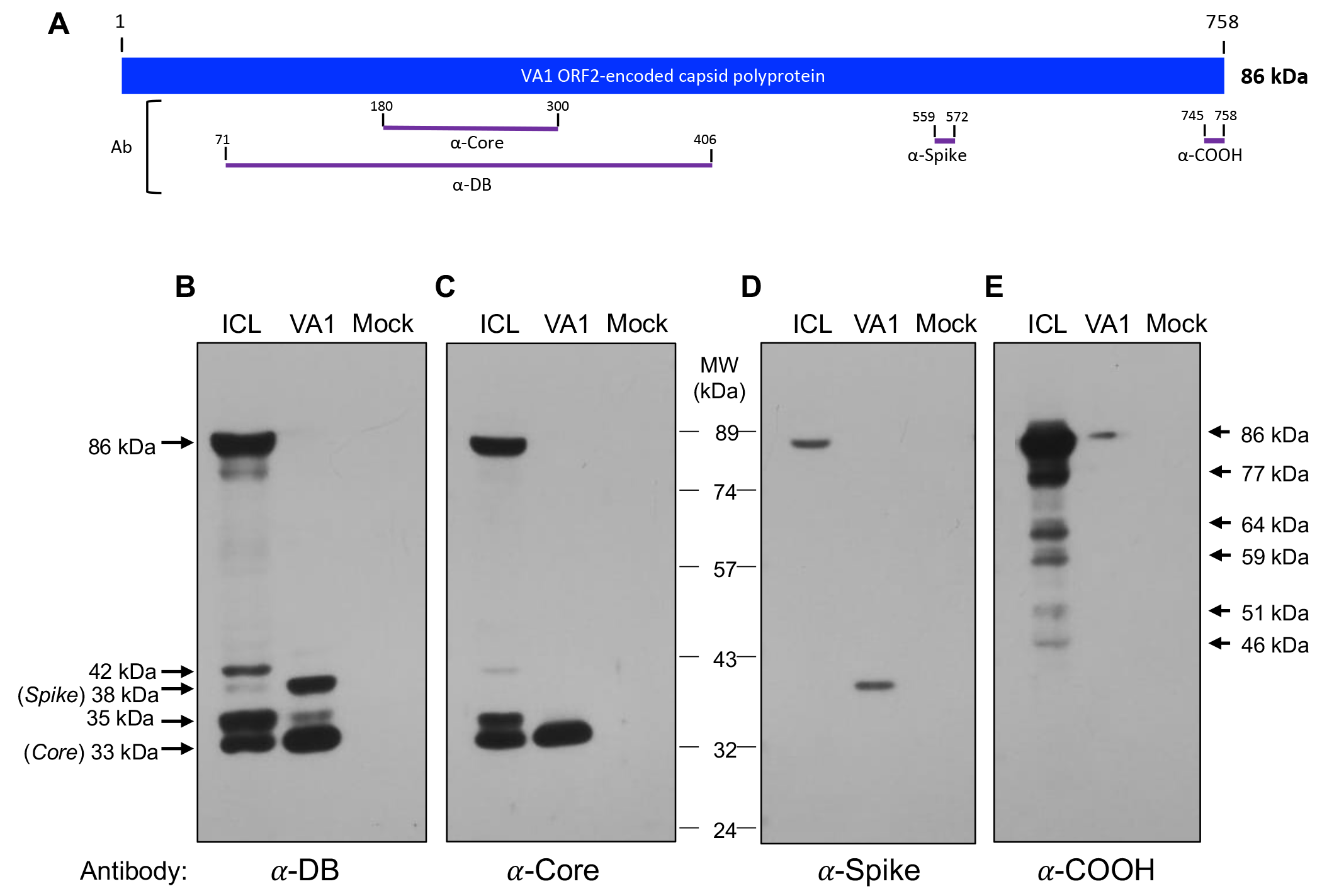
(A) Diagram of the VA1 ORF2 capsid precursor and recombinant proteins and peptides. ORF2 is represented as a box, and the recombinant proteins and the α-Spike and α-COOH peptides used to generate antibodies are represented as purple lines. The diagram is to scale, and the relative positions of the astrovirus VA1 recombinant proteins Core and DB, and the peptides Spike and COOH are shown; the numbers indicate the VA1 ORF2 amino acid residues included in each protein or peptide. (B-E) Identification of the core and spike capsid proteins assembled in the mature, infectious virus. Monolayers of Caco-2 cells were mock-infected or infected with VA1 at an MOI of 5, and the infection was left to proceed for 48 h at 37°C. At the end of the incubation period the cell media was discarded and the cell monolayers were washed twice with DMEM-HG and lysed by three cycles of freeze-thawing. Laemmli sample buffer was added, and the proteins were separated by 10% SDS-PAGE and analyzed by immunoblot using either (B) α-DB, (C) α-core, (D) α-spike, or (E) α-COOH antibodies. ICL, infected cell lysates; VA1, purified virus particles; Mock, mock infected cells. The MW markers are indicated between panels (C) and (D). The arrows at the left of panel (B) and at the right of panel (E) indicate the apparent molecular weight estimated for the viral proteins. The viral proteins of 33 and 38 kDa correspond to the core and spike domains of the mature, infectious virus (indicated).

The antibody directed against the spike region (α-spike) only recognized the precursor 86-kDa protein in the infected cell lysate, and a protein of 38 kDa in the purified virus (Fig. 3D), indicating that this protein corresponds to the mature spike protein. Finally, the antibody directed against the carboxy terminus region of the ORF2-encoded polyprotein (α-COOH) recognized the precursor protein and intermediate processing products of various molecular weights in the cell lysate, and it only recognized trace amounts of the precursor protein in the purified virus (Fig. 3E), showing that the 86 kDa protein is not present in the mature virus, or only in very minor proportion. These results indicate that the original DB antibody indeed recognizes the core protein, but also a portion of the spike protein. No protein bands were observed in uninfected cell lysates with any of the four anti-ORF2 protein antibodies evaluated (Fig. 3).

### Identification of the spike domain amino terminus

To determine the cleavage site that produces the core and spike proteins, virus particles were purified as described in Material and Methods, and the viral proteins were separated in an 11% SDS-polyacrylamide gel and stained with Coomassie blue. Only two major bands of 33 (VP33, core) and 38 kDa (VP38, spike) were observed (data not shown). These bands were excised from the gel and subjected to amino-terminal sequencing by Edman’s degradation. The amino terminus of VP33 did not yield a sequence, suggesting that the end of the core protein is blocked. On the other hand, the band corresponding to the VP38 protein rendered the sequence TDVQY, which corresponds to amino acid residues 348 to 352 of the VA1 ORF2-encoded polyprotein (see Fig. 6A. These results confirm the identification of the core and spike proteins present in the mature, infectious virus.

### VA1 replication does not depend on caspase cleavage of the capsid precursor

Classical astroviruses require that the intracellular 90 kDa capsid precursor, assembled in immature VP90 particles, is cleaved by caspases to VP70 for the egress of the VP70-particles to the extracellular medium (7). Thus, we explored whether this class of proteases could be involved in the intracellular proteolytic cleavage of the VA1 capsid polyprotein precursor. For this, Caco-2 cells were infected with VA1 at an MOI of 3 and incubated for 48 h in the presence of 50 μM of the pan-caspase inhibitor Z-VAD-FMK, and the processing of the VA1 precursor protein was assessed by Western blot using the DB antibody. The cleavage pattern of VA1 was the same, regardless the presence or not of the caspase inhibitor (Fig. 4A). As positive control, Caco-2 cells were infected with HAstV-8 in the absence or presence of Z-VAD-FMK. In the absence of the inhibitor, the VP90 and VP70 forms were observed (14) (Fig. 4B), indicating the cleavage of the polyprotein precursor. On the other hand, in the presence of the inhibitor only the VP90 protein was detected, an indication that the VP90 to VP70 cleavage was prevented, as previously reported (14, 15). In addition to the results described above, we found that the production of infectious VA1 virus was not affected by the caspase inhibitor Z-VAD-FMK (Fig. 4C), confirming that the VA1 astrovirus capsid precursor is not processed by caspases.

**Fig. 4.**
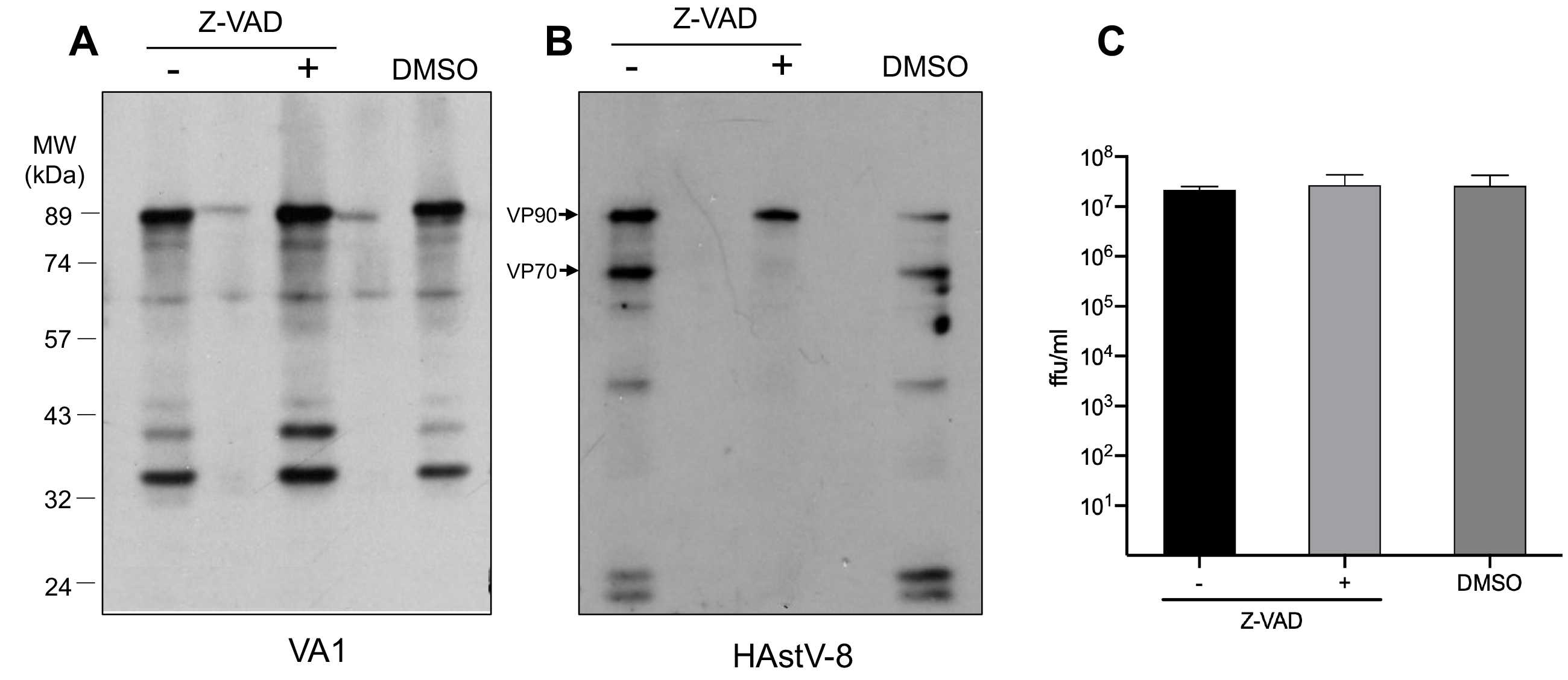
Caspases are not involved in processing the VA1 capsid precursor. Monolayers of Caco-2 cells were infected with either VA1 or HAstV-8 (strain Yuc8; previously activated by incubation with 200 μg/ml of trypsin for 1h at 37°C) at an MOI of 3. For this, the virus was adsorbed for 1 h at 37°C and then washed twice with MEM, and DMEM-HG (Z-VAD minus), DMEM-HG containing 50 μM of Z-VAD-FMK (Z-VAD plus), or DMEM-HG with the equivalent concentration of DMSO (DMSO) as a control, was added and the cells were incubated for 48 hat 37°C. After this time the cell media were discarded and the viral proteins in the cellular fraction were separated by 10% SDS-PAGE and analyzed by Western blot using (A) α-DB antibody to detect VA1, or (B) a rabbit polyclonal antibody to HAstV-8. (C) In parallel samples, the infectivity for astrovirus VA1 was determined by an immunoperoxidase focus-forming assay as described in Material and Methods. Data represent the mean±SD of three independent experiments.

### The VA1 capsid precursor is sensitive to trypsin cleavage

As stated above, unlike classical astroviruses, astrovirus VA1 does not need to be treated with trypsin to activate its infectivity. However, whether trypsin could cleave the VA1 capsid proteins and/or activate VA1 infectivity has not been analyzed. To evaluate this possibility, capsid proteins in the pellet and in the supernatant obtained during the virus purification method after centrifugation through a sucrose cushion, and prior to the CsCl gradient (see Material and Methods), were treated with increasing amounts of trypsin for 1 h at 37 °C and analyzed by Western blot using the DB antibody, and the viral infectious titer was also determined (Fig. 5).

**Fig. 5.**
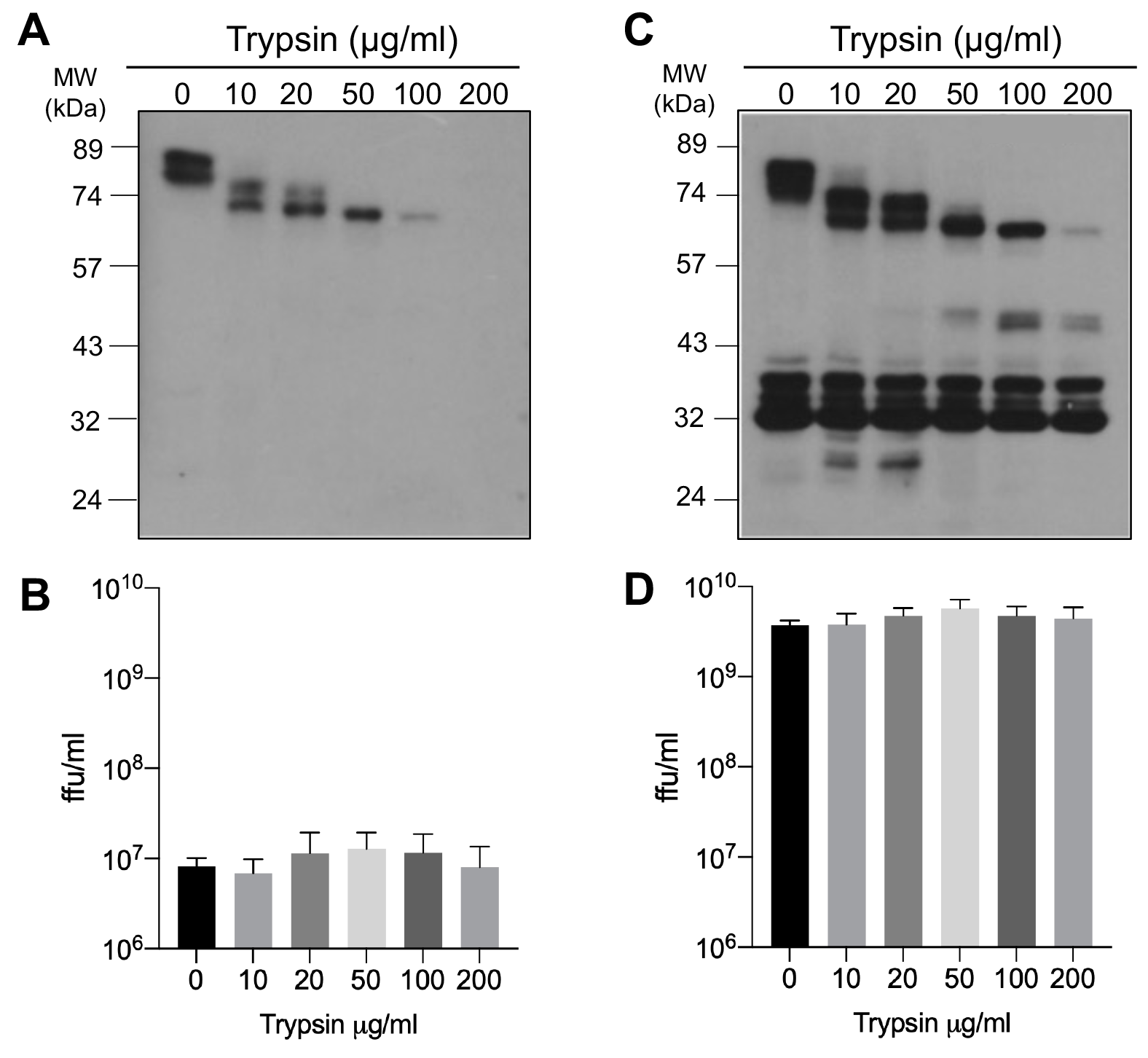
Trypsin cleavage of the VA1 capsid precursor does not lead to infectious virus. Monolayers of Caco-2 cells were infected with VA1 at an MOI of 5, and the infection was left to proceed for 48 h at 37°C. The cells were then scrapped into the cell medium and processed for virus purification as described in Material and Methods; after centrifugation through a 1 ml 30% sucrose cushion in TNE buffer, the supernatant was collected and the pellet was resuspended in TNE buffer. Aliquots of both supernatant (A) and pellet (C) fractions were treated with the indicated concentrations of trypsin during 1 h at 37°C; the proteins were separated by 10% SDS-PAGE and analyzed by Western blot using the α-DB serum. The MW markers are indicated. Parallel aliquots of both supernatant (B) and pellet (D) fractions were used to determine the virus infectivity titer by an immunoperoxidase focus-forming assay as described in Material and Methods. Data represent the mean±SD of three independent experiments.

**FIG. 6.**
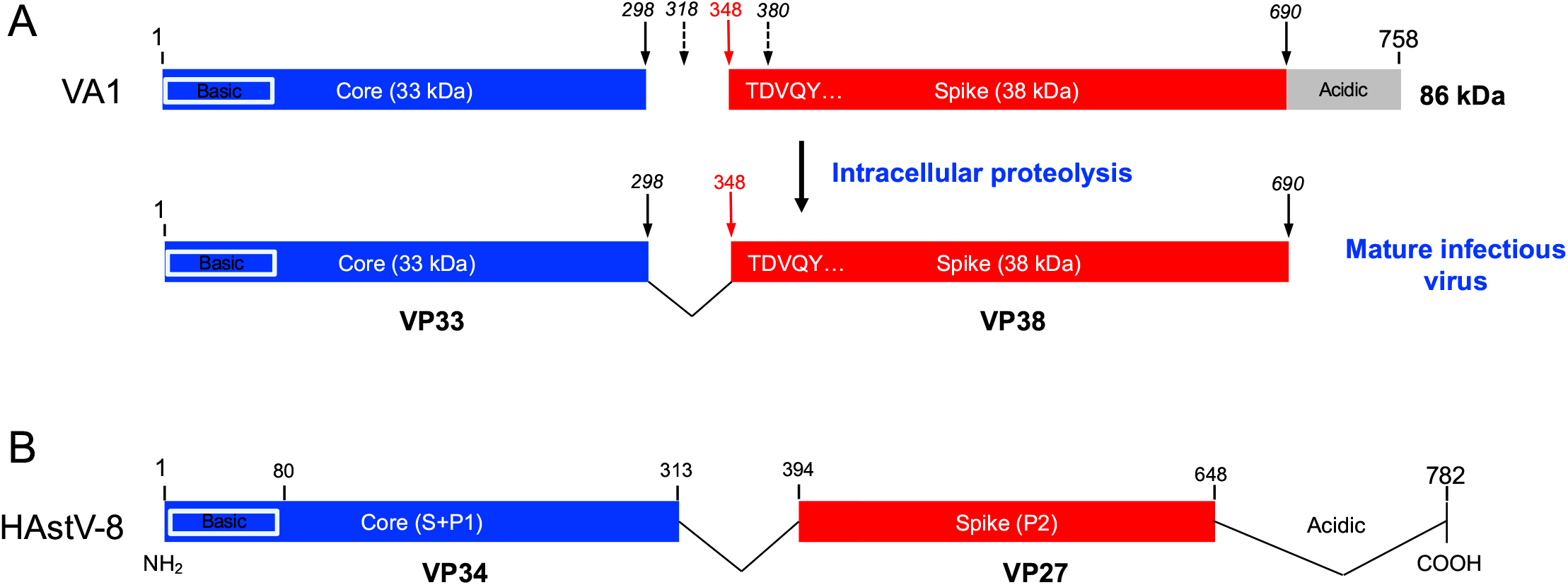
(A) Proposed proteolytical processing for the VA1 ORF2 capsid precursor. The blue and red boxes represent the protein products observed during virus replication; the stripped box represent a connecting peptide that is probably lost during the maturation process of the precursor. The basic and acidic domains of the precursor are indicated. The red arrow indicates the amino terminus of the spike domain VP38 determined by N-terminal sequencing of the protein; the continuous black arrows represent the estimated carboxy terminus of VP33 and VP38, based on the calculated apparent molecular weights of the proteins that constitute the mature, infectious virus; the dashed arrows represent cleavages that produce intermediate proteolytical products that produce core proteins precursors VP36 and VP42. (B) Trypsin cleavage products that constitute the mature, infectious human serotype 8 classical virus.

The main protein detected in the supernatant (the material that did not go through the sucrose cushion) was the 86-kDa capsid precursor, which was completely degraded upon treatment with a concentration of 200 μg/ml of trypsin (Fig. 5A). Of note, we detected a small amount of infectivity in these samples, treated or not with trypsin (Fig.5B), suggesting that small amounts of infectious virus were present in the supernatant fraction, and that its infectivity was not affected by trypsin. In contrast, in the pelleted sample that went through the sucrose cushion, the 33-, 35- and 38-kDa proteins were enriched, although the 86-kDa precursor protein was also detected in the non-treated sample (Fig. 5C). Of interest, when this sample was treated with different concentrations of trypsin, the 86-kDa precursor protein was gradually degraded, as it had been seen in the supernatant sample, but the protease treatment did not affect the lower molecular weight proteins (Fig. 5C) and, similarly, the virus infectivity remained unchanged upon trypsin treatment (Fig. 5D). Altogether, these results suggest that the VA1 capsid proteins incorporated into mature, infectious virus are no longer sensitive to trypsin cleavage. In contrast, the 86-kDa precursor is either loosely assembled into virus particles and sensitive to trypsin degradation both in the supernatant and pellet fractions, or it is in a soluble form that was present in the supernatant and contaminated the sedimented virus. These findings also suggest that the infectious form of the virus is formed by the processed polypeptides and not by the precursor polyprotein.

## DISCUSSION

In this work we confirmed that, unlike classical human astroviruses, the neurotropic astrovirus strain VA1 does not need to be treated with trypsin to activate its infectivity. However, we showed that the ORF2-encoded polyprotein is indeed proteolytically processed during virus replication and that this process is required to generate infectious virus. The fact that the polyprotein precursor is cleaved during the virus replication cycle even in the presence of FCS, together with the observation that the cell-associated virus is also proteolytically cleaved in the absence of trypsin in the culture medium (Fig. 1), strongly supports the idea that the cleavage of the ORF2 precursor protein occurs intracellularly. Accordingly, the capsid protein of the infectious virus particles released into the cell medium is already processed (Fig. 1C).

Using antibodies directed to different regions of the ORF2-encoded polyprotein we found that mature viruses are composed by two proteins, one of 33 kDa (VP33) that represents the core protein of the virus, and a second of 38 kDa (VP38) that constitutes the spike protein of the virus particle (Fig. 3). In addition, we detected several cleavage intermediate products of the ORF2 polyprotein during the cleavage pathway that yields VP33 and VP38. The intermediate products detected varied from experiment to experiment and were also dependent on the antibody used. For instance, when antibodies directed to the carboxy terminus of the polyprotein precursor were used, in addition to the 86-kDa precursor protein (VP86), bands ranging from about 46 to 77 kDa were observed. This indicates that the ORF2 capsid protein can be shortened by cleavage from the amino terminus of the precursor to render at least 5 intermediate products (Fig. 3D). On the other hand, when antibodies to the core were used, it appears that processing of VP86 can also start at the carboxy-terminal region, since intermediate products of 35 and 42 kDa were observed (Fig. 3B). Thus, there seems to be alternative cleavage pathways that finally converge to render VP33 and VP38, the two polypeptides that constitute the mature infectious virus. The 35-kDa core intermediate product was always very abundantly observed in infected cells, although it was not detected in the mature virus. The expected molecular weight of the precursor protein, based on the amino acid composition of ORF2 that encodes a polypeptide of 758 amino acids is 84.2 kDa, however the apparent MW observed is 86 kDa. Whether this is due to atypical migration of the band during electrophoresis or post-translational modifications to the capsid protein is not clear.

A diagram showing the expected final products of the proteolytic cleavage pathway is shown in Fig. 6A, as well as the various potential protease cleavage sites deduced from the molecular weight of the core intermediate products. Considering the molecular weight of VP33 and the identified amino terminus of the spike, it seems that a segment of about 50 amino acids, is lost from the precursor protein, as has also been observed for the processing of the VP70 protein of HAstV-8 (11, 16). Our results, along with structural predictions, also suggest that the N-terminal region of the VA1 spike protein contains one or more beta strands between residues 350-385 that tether it to the core protein, like the N-terminal region of the spike of classical human astroviruses (17). Of interest, the spike protein of VA1 is larger than the core protein, as opposed to that observed in classical human astroviruses (7). It will be very interesting to determine the structures of these proteins as well as the VA1 virus particle to understand the structural basis for these differences.

In contrast to what has been previously shown to occur for HAstV-8, caspases were not found to be involved in the processing of the VA1 VP86 precursor. On the other hand, and as expected, trypsin treatment did not affect the VP33 and VP38 proteins once they were assembled into VA1 virions (Fig. 5), given that the virus infects in a trypsin-like environment in the human gut. However, the precursor protein and most of the intermediate products were readily digested by trypsin, suggesting that the non-particulated precursor protein is sensitive to this protease. Alternatively, it is possible that assembled VP86 is susceptible to trypsin cleavage before it is processed into VP33 and VP38. Furthermore, the observation that the infectivity of the virus did not decrease despite of the precursor protein being completely degraded after trypsin treatment, indicates that the infectious virus form is constituted by the processed core and spike polypeptides.

Whether the protease involved in the cleavage of VP86 is of cellular or viral origin remains to be determined, although the participation of more than one protease is also possible. We are currently investigating the origin of the protease(s), but if a viral protease is involved, it will represent a major difference with classical astroviruses. Also, the fact that this virus has been reported to infect and replicate in neural cells and to be associated with encephalitis/meningitis in immunosuppressed individuals is consistent with its independence of trypsin activation *in vivo*; these observations are also consistent with the involvement of an intracellular or viral protease in the process.

In this work we also report a single-cycle replication curve of VA1, assessing the production of infectious virus associated to cells and released into the cell medium up to 72 hpi. The replication cycle seems to be completed in 48 h, with an eclipse period of about 12 h. Remarkably, no cytopathic effect was observed. These observations are similar to those previously reported for VA1 multistep growth curves carried out in CaCo-2 cells (13, 18), in which genome viral copies rather than infectious virus were assessed. Also, similar growth curves were reported in HEK293T and A549 cells (13), human intestinal enteroids (19), and neural cells (18). These observations contrast with the replication cycle of HAstV-8 in Caco-2 cells, which has an eclipse period of about 6 to 8 h and the synthesis of infectious viruses plateau at 24 hpi ((16), unpublished data); these observations suggest a more efficient replication cycle of classical viruses in human cells. The findings reported in this work advance our knowledge about the biology of the neurotropic VA1 human astrovirus and set the basis to better understand the process of virus entry and the proteolytical cleavage of viral polyproteins.

## Material and Methods

### Virus and cells

Cells from a human colon adenocarcinoma (Caco-2), clone C2Bbe, obtained from the American Type Culture Collection (ATCC) were used in this work. Cells were grown in DMEM [Dulbecco’s modified Eagle’s medium-reduced serum (DMEM-RS; Thermo Scientific HyClone, Logan, UT, USA)] supplemented with 5% heat-inactivated fetal calf serum (FCS; Biowest, Kansas City, MO, USA) at 37°C in a 10% CO_2_ atmosphere. Astrovirus VA1 (AstV-VA1) was kindly provided by David Wang (Washington University) and was propagated in Caco-2 cells. The virus, at a multiplicity of infection (MOI) of 0.01-0.001, was adsorbed to the cells for 1 h at 37°C. The unbound virus was removed, and the cells were incubated for four days at 37°C in Dulbecco Modified Essential Medium (DMEM) (Sigma Cat. D7777), supplemented with nonessential amino acids (Gibco Cat. 11140) without serum. Finally, the cells were washed twice with DMEM-HG and lysed by three cycles of freeze-thawing. The resulting infected cell lysate was aliquoted and stored at -70°C. Astrovirus serotype 8 (HAstV-8) at an MOI of 0.5 was adsorbed to Caco-2 cells; the virus inoculum was previously activated with trypsin (200 μg/ml, Gibco Cat. 27250-18) for 1 h at 37 °C, and soybean trypsin inhibitor (200 μg/ml) was added just before inoculation to the cells. After this time the unbound virus was removed, and MEM without serum was added. The infected cells were incubated at 37°C and harvested 3 days post-infection. Finally, the cells were washed twice with DMEM-HG and lysed by three cycles of freeze-thawing. The resulting viral lysate was aliquoted and stored at -70°C.

### Antibodies

A protein containing amino acid residues 71 to 403 of VA1 ORF2 (GenBank accession number NC_013060.1) was synthesized in *E. coli* and purified as described (20), and was used to generate a hyperimmune serum in New Zealand rabbits, as previously reported (21) (DB antibody). Similarly, a polypeptide containing amino acid residues 180 to 300 of VA1 ORF2 was produced in *E. coli* as a fusion with glutathione S-transferase (GST) using the pGEX-4T-1 vector (Pharmacia). A hyperimmune rabbit serum was produced as above (α-core antibody). Two synthetic peptides and their corresponding rabbit anti-antibodies were obtained from GeneScript. These peptides represent ORF2 amino acids 559-VLSEPEDFDVVQKS-572 (α-spike antibody) and 745-AIAVKKKLRRGHAE-758 (α-COOH antibody). Both antibodies had an extra-cysteine added at the amino terminus. Antibodies to HAstV-8, strain Yuc8, have been reported (11).

### Virus purification

VA1 virus was purified as reported (8, 12), with some modifications. Briefly, confluent monolayers of Caco-2 cells were infected with VA1 at an MOI of 5, and the cells were incubated for 48 h at 37°C. The cells were then scraped into the cell medium, and centrifuged at 2000×*g* for 10 min at 4°C. The supernatant was discarded and the pellet was resuspended in 1% Triton X-100 (Sigma) in TNE buffer (50 mM Tris-HCl, pH 7.4, 0.15 M NaCl, 10 mM EDTA) and incubated for 30 min on ice with frequent stirring. After centrifugation at 2000×*g* for 10 min at 4°C, the supernatant was collected and ultracentrifuged at 50,000×*g* for 16 h at 4°C. The resulting pellet was resuspended in 4 ml of TNE buffer and sonicated to better solubilize the pellet. This suspension was centrifuged through a 1 ml 30% sucrose cushion in TNE buffer at 200,000×*g* for 2 h at 4°C. The virus in the pellet was then purified by CsCl density centrifugation. For this, the pellet was resuspended in 4 ml TNE, sonicated, and 2.4 g of CsCl were added. After centrifugation at 160,000x*g* for 18 h at 4°C the opalescent band corresponding to viral particles was collected, diluted to a 5 ml final volume with TNE and centrifuged at 200,000 ×g for 2h at 4 °C. The resulting pellet was resuspended in 400 μl TNE.

### Infectivity assay

Caco-2 cells were grown to confluence in 96-well plates. The cells were infected with 50 μl of 2-fold serial dilutions of the viral lysate for 1 h at 37°C. Then, the non-adsorbed virus was removed, and the cells were incubated at 37°C for 48 h. Afterward, the cell monolayers were processed for an immunoperoxidase assay. Briefly, the cell monolayers were fixed with 2% formaldehyde in PBS for 20 min at room temperature and then washed twice with PBS. The cell membrane was then permeabilized with 0.2% Triton X-100 (Sigma) in PBS for 15 min at room temperature. Next, samples were immunostained using a monoclonal neutralizing antibody (2A2, ref (5)), followed by incubation with protein A coupled to peroxidase. Finally, the cells were washed twice with PBS and stained with 50 μl of a solution containing 1 mg/ml of 3-amino-9-ethyl carbazole (AEC), 0.04% H_2_O_2_, and sodium acetate buffer (50 mM, pH 5). The reaction was stopped by washing in tap water. The number of focus forming units (FFU) was determined in a Nikon TMS inverted phase-contrast microscope with a 20x objective. The virus titer was expressed as FFU/ml.

### Western blots

Cell monolayers grown to confluence in 48-well plates were infected at an MOI of 3 or 5, depending on the experiment. After 48 h incubation at 37°C, cells were lysed in Laemmli sample buffer. The proteins were separated in a 10% SDS-polyacrylamide gel and transferred to Immobilon NC membranes (Millipore Merck, Darmstadt, Germany). The membranes were blocked with 5% non-fat dry milk in PBS for 1 h at room temperature and then incubated with primary antibodies (DB 1:2000, α-core 1:2000. α-spike 1:2000, α-COOH 1:2000, α-HAstV-8 1:2000) diluted in PBS containing 0.1% non-fat dry milk. The membranes were then incubated with anti-IgG rabbit antibodies conjugated to horseradish peroxidase (PerkinElmer Life Sciences (Boston, MA, USA), and developed using the Western Lightning Chemiluminescence Reagent Plus (PerkinElmer). To determine the apparent molecular weight (MW) of the proteins in the developed films, we used the PageRuler prestained protein ladder (Product#26616, Thermo Scientific). However, since, the proteins in this ladder are bound to chromophores, their MW are only approximate, and vary by about 5% from lot-to-lot. Thus, we recalibrated the MW of these markers with the PageRuler unstained protein ladder (Product#26614, Thermo Scientific) for a more precise MW determination. The apparent MW shown in the Western blot images correspond to the calibrated MW.

### ZVAD and trypsin treatments

Caco-2 cells grown in a 48-well plate were infected with either VA1 or Yuc8 at an MOI of 3 (a MOI of 5 was used for the VA1 trypsin treatment experiment). The virus was adsorbed for 1 h at 37°C and then washed twice with MEM. Cells were treated with DMEM-HG containing 50 μM of Z-VAD-FMK (627610; Sigma-Aldrich, St. Louis, MO, USA), or DMEM-HG with the equivalent concentration of DMSO as a control. For both viruses, the cells were incubated for 48 h at 37°C and then frozen and thawed 3 times. The infectivity of astrovirus VA1 was determined by an immunoperoxidase assay, and the viral protein production of both viruses was analyzed by Western blot, using the α-DB serum to VA1, or a rabbit polyclonal antibody to HAstV-8. For the trypsin-treatment assays, the supernatant and pellet obtained after virus centrifugation through a sucrose cushion (see above) were treated with different trypsin concentrations (0, 10, 20, 50, 100 and 200 μg/ml) (Gibco, Life Technologies, Carlsbad, CA, USA) for 1 h at 37°C. Each treated sample was split into two fractions. One fraction was used to determine the viral infectivity immediately after trypsin treatment, and the proteins in the second fraction were separated in 10% SDS-polyacrylamide gels and analyzed for Western blot.

### N-terminal protein sequencing

CsCl-purified virus particles, were separated by SDS-PAGE and transferred to a polyvinylidene fluoride membrane (PVDF-Plus 0.1 μm; Osmonics) for 1 h at 50 V in CAPS [3-(cyclohexylamine)-1-propane sulfonic acid] buffer (10 mM CAPS [pH 11], 10% methanol). All reagents used were freshly prepared and filtered through a 0.22 μm pore-size membrane. Transferred proteins were visualized by staining with Ponceau-S, and the bands were excised, washed thoroughly with MilliQ water, and dried in a dust-free environment. Automatic amino acid sequencing determination, by Edman degradation was performed using a PPSQ-31A Protein Sequencer from Shimadzu Scientific Instruments, Inc. (Columbia, MD, USA), using the PVDF membrane with the transferred protein.

### Statistics

Statistical analysis was determined by a two-tailed T-test with confidence interval of 99%, using the GraphPad Prism 9.0.1 Software (GraphPad Software, Inc.).

## Acknowledgments

This research was partially supported by grants NIH R01 AI144090 to R.M.D. and C.F.A., CONACyT M0037-Fordecyt 302965 to S. López; and DGAPA IN210120 to T. López. We are grateful to David Wang (Washington University) for kindly providing human astrovirus VA1. We thank Rafaela Espinosa for her advice and help in cell culture, and we also thank E. Mata and the animal house personnel for help during animal handling and immunization.

